# Identifying Meta-QTLs for Stay-Green in Sorghum

**DOI:** 10.1101/2021.08.06.455413

**Authors:** Ahmed Aquib, Shadma Nafis

## Abstract

To develop resilient crops it is necessary to understand the underlying genetics of climatic response. A strong association between stay-green and post-flowering drought tolerance in Sorghum has been established. Being a complex quantitative trait, Quantitative Trait Loci (QTL) mapping experiments of stay-green in Sorghum have been frequently performed. The objective of the current study was to find consensus genomic regions that control stay-green by integrating the QTLs mapped in previous studies. Meta-QTL analysis was performed to summarize 115 QTLs projected on the consensus map. The analysis generated 32 Meta-QTL regions within which candidate gene (CG) identification was undertaken. 7 candidate genes were identified using the markers tightly linked to the Meta-QTLs. The results from this study will facilitate future attempts aiming to improve and understand drought tolerance in Sorghum.

## Introduction

Sorghum [Sorghum bicolor (L.) Moench] is an important staple and fodder crop, especially for the people living in semi-arid regions of Asia and Africa. Its resilience to osmotic stress has made it an exceptional model for studying genomics for drought tolerance. Stay-green trait, which is the ability to resist premature senescence in response to post flowering drought, has been recognised as one of the most crucial drought related traits studied in Sorghum [1]. Stay-green genotypes in sorghum are able to retain photosynthetically active green leaf area and normal grain filling under water limited conditions [2],[3],[4]. The underlying genetic factors affecting stay-green, a quantitative trait, have been extensively studied through QTL mapping experiments. Over 100 QTL for stay-green traits in Sorghum have been mapped in different populations using a different set of markers.

Differences in population, environment and choice of markers limit the transferability of QTL across breeding programs. Also, the genetic effects of QTLs identified in any single study needs to be tested for their presence in diverse genetic backgrounds and environments. QTL meta analysis is an approach to identify consensus genomic regions by combining QTLs from diverse populations and environments [5]. These consensus QTL, also called MetaQTL (MQTL), are stable and robust regions where QTLs were mapped recurrently in experiments. Additionally, meta-QTL analysis leads to a significant reduction in confidence interval in the MQTL compared to the original QTLs. The reduced interval and reliability of MQTLs over QTLs enable Candidate Gene (CG) identification and Marker Assisted Selection (MAS) with improved precision.

In this study, we have performed the first meta-QTL analysis for stay-green traits in Sorghum using results from 9 previous QTL mapping experiments Table 1. The analysis involved construction of consensus maps, QTL projection and statistical clustering of QTLs. Thereafter, an attempt was made to identify candidate genes (CGs) in the MQTL regions. The results from this meta analysis can be useful in improving sorghum for drought tolerance. The identified CGs provide a framework for a better understanding of underlying genetic mechanisms for stay-green and its relation with osmotic stress tolerance.

**Table 1.** Summary of QTL association studies considered for the Meta-QTL analysis

| S.No. | Experiment Name | Number of Stay-green QTL(s) | Parents of population | Population type | Population size |
| --- | --- | --- | --- | --- | --- |
| 1 | Crasta et al. 1999 | 7 | B35 x Tx430 | RIL | 96 |
| 2 | Hausmann et al. 2002 (RIP-1) | 19 | IS9830 x E36-1 | RIL | 226 |
| 3 | Hausmann et al. 2002 (RIP-2) | 21 | N13 x E36-1 | RIL | 226 |
| 4 | Xu et al. 2000 | 3 | B35 x Tx7000 | RIL | 98 |
| 5 | Kebede et al. 2000 | 14 | SC56xTx7000. | RIL | 125 |
| 6 | Sabadin et al. 2012 | 4 | BR007 x SC283 | RIL | 100 |
| 7 | Srinivas et al 2008 | 9 | 296B x IS18551 | RIL | 168 |
| 8 | Subudhi et al. 2000 | 4 | B35 x Tx7000 | RIL | 98 |
| 9 | Tau et al. 1999 | 5 | QL39/QL41 | RIL | 152 |
| 10 | Reddy et al 2014 | 43 | M35-1 X B35 | RIL | 245 |

## Materials and Methods

### Collection of Stay-Green QTLs in Sorghum

Through a rigorous literature search, a database of Sorghum stay-green QTLs was created on a spreadsheet. The database consisted of 130 QTLs published across 9 studies from 1999 to 2014. The columns of the spreadsheet contained for each QTL the following information: (1) Name, (2) Chromosome, (3) Linkage group, (4) Phenotypic variance explained or R2, (5) Position, (6) 95 Percent Confidence Interval and (7) Source. For each QTL a 95 percent confidence interval was calculated using the formula 163/NR2 [15]. 4 of the QTLs which had confidence intervals greater than 50cM were screened from further analysis. Additionally data about each of the 9 studies was collected and saved separately as metadata. The metadata consisted of the name, population type, population size, mapping function and cross performed in each study.

### Consensus Map Building

In order to study relative positions of the range of QTLs mapped with different types of markers, a consensus map was built which contained markers from all the experiments. Apart from the maps described in the studies that reported QTLs, two Sorghum consensus maps published on Gramene database were also used for building the consensus map [16], [17], [18]. In some of the QTL association experiments, such linkage maps were used where linkage groups were arbitrarily named. To such linkage groups size-based chromosomal nomenclature (SBI-01 to SBI-10) was applied. Link-age groups were assigned the chromosome with which the maximum number of markers matched with the help of an R script. The ConsMap tool originating from the MetaQTL package and implemented in Biomercator V4.2 was used to compile all the maps into a single consensus map [19], [20].

### QTL Projection and Meta Analysis

Using the QTLproj tool in Biomercator V4.2, adapted from the MetaQTL package, 115 out of 126 QTLs were projected on the consensus map developed in this study [19], [20]. QTLproj uses flanking markers on the original map and the marker interval distances in order to find the optimal position and confidence interval for the QTL on the consensus map. The QTLClust tool from the MetaQTL package was used to probabilistically cluster QTLs. The projected QTLs on the consensus map were used for the meta analysis. The clustering procedure for the algorithm is based on a gaussian distribution of the observed QTLs around their position with variance derived from the confidence interval. The best meta model was chosen according to the Akaike Information Criterion (AIC). The meta-QTLs that correspond to only a single QTL were not considered.

### Collection of the Physical Positions of the Markers

Anchors are genetic markers having their physical location on the genome mapped. The availability of anchors is necessary to approximate the physical location of genetic intervals such as QTLs or MQTLs. In this study we have combined the marker location data generated previously in two studies in order to locate MQTLs on the reference genome[21] [22]. In addition, 32 more markers were mapped on the Sorghum reference genome through sequence retrieval from databases (Gramene and NCBI) and a subsequent BLAST search [16], [23], [24].

### Locating MQTLs on the Reference Genome

Predicting a physical interval to the MQTLs is challenging due to the deviation of the relationship between genetic and physical distances from linear. Previously, studies have used two flanking markers around a MQTL to predict the physical intervals. But, using only two flanking markers for prediction does not take into account the variation in positions of markers within the MQTL region, or near to it. Therefore, in the current study, a linear regression model was developed for each MQTL region using anchors within MQTL regions and flanking it. If there were no anchors within the region then three nearest anchors were used. A 95 percent confidence level was used for prediction. Only the MQTLs with predicted physical intervals shorter than 10 Mb were used for candidate gene mining. Due to lack of anchors and therefore larger confidence intervals on the physical maps, some distinct MQTLs on the genetic map cannot be resolved separately on the reference genome, i.e their CI overlapped. A non-redundant region was then taken as the physical interval of all such MQTLs combined. Through this we were able to identify 7 non-redundant regions consisting of 1-4 MQTLs.

### Candidate Gene Mining

Gene sets from the 7 regions of interest were retrieved by querying and subsetting the v3.1.1 gene annotation of Sorghum bicolor, containing a total of 34,118 genes, from the Ensembl-Plants database [25], [26]. BiomaRt and GenomicRanges packages from Bioconductor were used for retrieving the gene sets using the R programming language [27], [28], [29], [30]. The genes retained after filtering the Gene Ontology term with the keywords “chlorophyll biosynthetic process” and “chlorophyll catabolic process” were considered as candidate genes [31], [32]. The information about the function of genes was acquired from uniprot knowledge-base by searching orthologous rice genes [33], [34].

## Results

### Distribution of QTLs and MQTLs

From the initial pool of 130 QTLs, 115 QTLs for stay-green trait from 9 studies were aligned on a single consensus map consisting of 10 chromosomes of Sorghum. The remaining QTLs were not projected either due to high CI (>50 cM) or couldn’t be projected due to lack of common markers. The meta-analysis was able to identify 32 Meta-QTL regions from the projected QTLs. A summary of chromosome wise distribution of these QTLs and MQTLs is shown in Fig 1. Chr 1 had the highest number of QTLs reported (N=26), followed by Chr 3 (N=23) and Chr 2 (N=19). Chr 3 harbored the maximum number of MQTLs per chromosome whereas Chr 6 harbored no MQTL. The average phenotypic variance explained (PVE) in all the QTLs was 10% with 97% QTLs having PVE over 5%. The average CI for the MQTLs was 6.1 cM, which corresponds to a 46% reduction from the average CI of original QTLs. The number of individual QTLs per MQTL ranged from 2 to 9. 26 MQTLs consisted of original QTLs discovered in distinct association experiments, hence could have greater stability across environments.

**Fig. 1.**
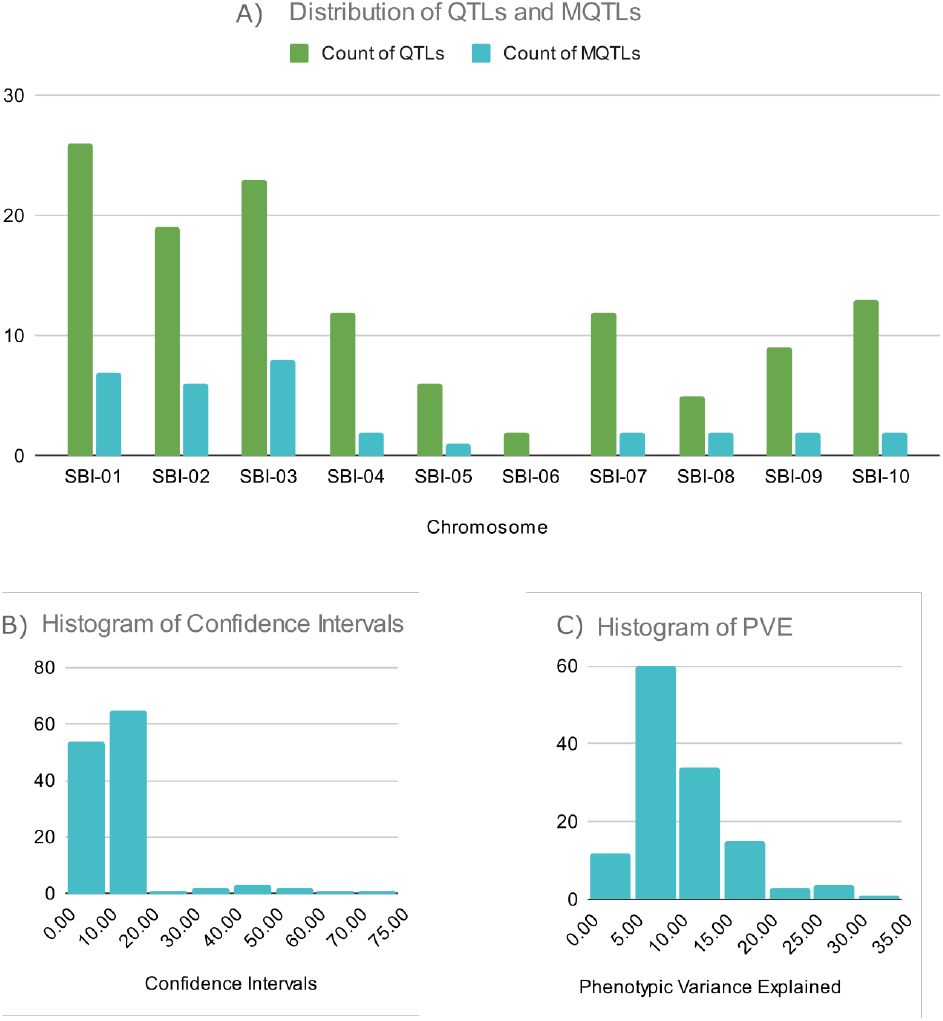
A) Chromosome wise distribution of the original QTL and MQTLs; B) A histogram of confidence intervals of the original QTLs; C) Histogram of the phenotypic variance explained (R^2^ of the original QTLs (in %))

Out of 1050 markers in the complete consensus map, only the physical positions of 232 ( 22%) were known on the reference genome. Due to lack of anchors the attempt to estimate the location of QTLs on the sequenced reference genome of Sorghum was only successful for 13 out of 32 meta QTLs. The criterion for a successful estimation was a prediction, with 95% confidence level, of an interval lower than 10 Mb. Furthermore, on the reference genome the predicted intervals of some of these MQTLs overlapped. Therefore for the purpose of candidate gene identification, the 13 MQTLs with known physical locations were grouped into 7 non-redundant regions.

### Candidate Gene Analysis

A total of 4587 genes were present within the 7 regions consisting of 13 MQTLs on chr 2, chr 3, chr 4 and chr 10. The interval sizes ranged from 3.24 to 13.38 Mb and the number of genes per region ranged from 290 to 1612. Of the total 4587 genes 7 genes related to chlorophyll biosynthesis and degradation were selected as candidate genes through filtering the Gene Ontology (Go Term). Region 3, 5, and 6 corresponding to MQTL_SG_14, MQTL_SG_21, and MQTL_SG_23 respectively, had no genes related with chlorophyll metabolism. 5 of the 7 genes function in the chlorophyll biosynthesis process and 2 in chlorophyll degradation. The genes involved in chlorophyll biosynthetic process were: (1) Polynucleotide phosphorylase 1, (2) Isopentenyl-diphosphate Delta-isomerase, (3) Uroporphyrinogen decarboxylase, (4) TP_methylase domain-containing protein, and (5) Delta-aminolevulinic acid dehydratase. Whereas the genes involved in degradation of chlorophyll were (1) Uncharacterized Protein (SORBI_3002G338100), and (2) AMP-binding domain-containing protein. Table 3 gives a tabular overview of the genes of interest within the non-redundant physical intervals containing MQTLs.

**Table 2.** Summary of all the Meta-QTLs generated

| MQTL | Chromosome | Position on the consensus map (cM) | Confidence interval (cM) | Nearest Marker | N QTLs | N studies | Position on the Sorghum genome |
| --- | --- | --- | --- | --- | --- | --- | --- |
| MQTL_SG_1 | SBI-1 | 31.29 | 4.99 | umc27 | 4 | 3 | - |
| MQTL_SG_2 | SBI-1 | 46.89 | 3.12 | umc128 | 3 | 1 | - |
| MQTL_SG_3 | SBI-1 | 71.81 | 4.23 | txs645 | 9 | 3 | - |
| MQTL_SG_4 | SBI-1 | 82.65 | 7.69 | txp88 | 4 | 2 | - |
| MQTL_SG_5 | SBI-1 | 94.95 | 9.72 | txp32 | 3 | 2 | - |
| MQTL_SG_6 | SBI-1 | 129.78 | 8.9 | isp237 | 4 | 2 | - |
| MQTL_SG_7 | SBI-1 | 182.71 | 6.41 | st1916 | 3 | 2 | - |
| MQTL_SG_8 | SBI-2 | 20.55 | 8.95 | csu13 | 2 | 2 | 3.227 - 6.956 |
| MQTL_SG_9 | SBI-2 | 115.13 | 8.01 | X13/61-259 | 5 | 4 | 57.858 - 65.883 |
| MQTL_SG_10 | SBI-2 | 123.26 | 6.09 | txp179 | 6 | 4 | 60.282 - 67.847 |
| MQTL_SG_11 | SBI-2 | 141.39 | 5.69 | umc4 | 2 | 1 | 65.415 - 71.063 |
| MQTL_SG_12 | SBI-2 | 155.23 | 3.79 | psr107 | 6 | 1 | 69.912 - 71.24 |
| MQTL_SG_13 | SBI-2 | 162.91 | 5.25 | Bnl8.37 | 3 | 1 | - |
| MQTL_SG_14 | SBI-3 | 35.36 | 5.21 | txp489 | 2 | 2 | 4.24 - 7.484 |
| MQTL_SG_15 | SBI-3 | 69.05 | 4.78 | spb9076 | 5 | 2 | - |
| MQTL_SG_16 | SBI-3 | 83.64 | 3.96 | rz323.2 | 6 | 2 | 51.566 - 53.941 |
| MQTL_SG_17 | SBI-3 | 90.4 | 3.46 | msbcir276 | 8 | 2 | 51.705 - 58.809 |
| MQTL_SG_18 | SBI-3 | 106.97 | 7.51 | txp2 | 5 | 4 | 56.076 - 57.899 |
| MQTL_SG_19 | SBI-3 | 121.63 | 6.46 | wg889 | 5 | 5 | 57.605 - 59.632 |
| MQTL_SG_20 | SBI-3 | 132.01 | 2.91 | spb9139 | 5 | 3 | - |
| MQTL_SG_21 | SBI-3 | 147.67 | 7.95 | rubp | 2 | 1 | 62.478 - 66.593 |
| MQTL_SG_22 | SBI-4 | 43.25 | 9.27 | txp26 | 4 | 3 | - |
| MQTL_SG_23 | SBI-4 | 111.19 | 2.32 | spb9303 | 5 | 2 | 50.51 - 53.857 |
| MQTL_SG_24 | SBI-5 | 48.59 | 2.29 | tx387 | 5 | 3 | - |
| MQTL_SG_25 | SBI-7 | 60.61 | 3.43 | isu38 | 5 | 2 | - |
| MQTL_SG_26 | SBI-7 | 147.62 | 2.05 | txp295 | 4 | 2 | - |
| MQTL_SG_27 | SBI-8 | 71.12 | 10.45 | spb4432 | 3 | 2 | - |
| MQTL_SG_28 | SBI-8 | 131.66 | 7.02 | X14/48-64 | 3 | 2 | - |
| MQTL_SG_29 | SBI-9 | 83.41 | 11.44 | st368 | 2 | 2 | - |
| MQTL_SG_30 | SBI-9 | 126.31 | 0.62 | gpsb079 | 7 | 1 | - |
| MQTL_SG_31 | SBI-10 | 112.58 | 9.33 | spb7058 | 3 | 2 | - |
| MQTL_SG_32 | SBI-10 | 127.06 | 12.53 | cup07 | 2 | 2 | 56.742 - 62.441 |

**Table 3.**
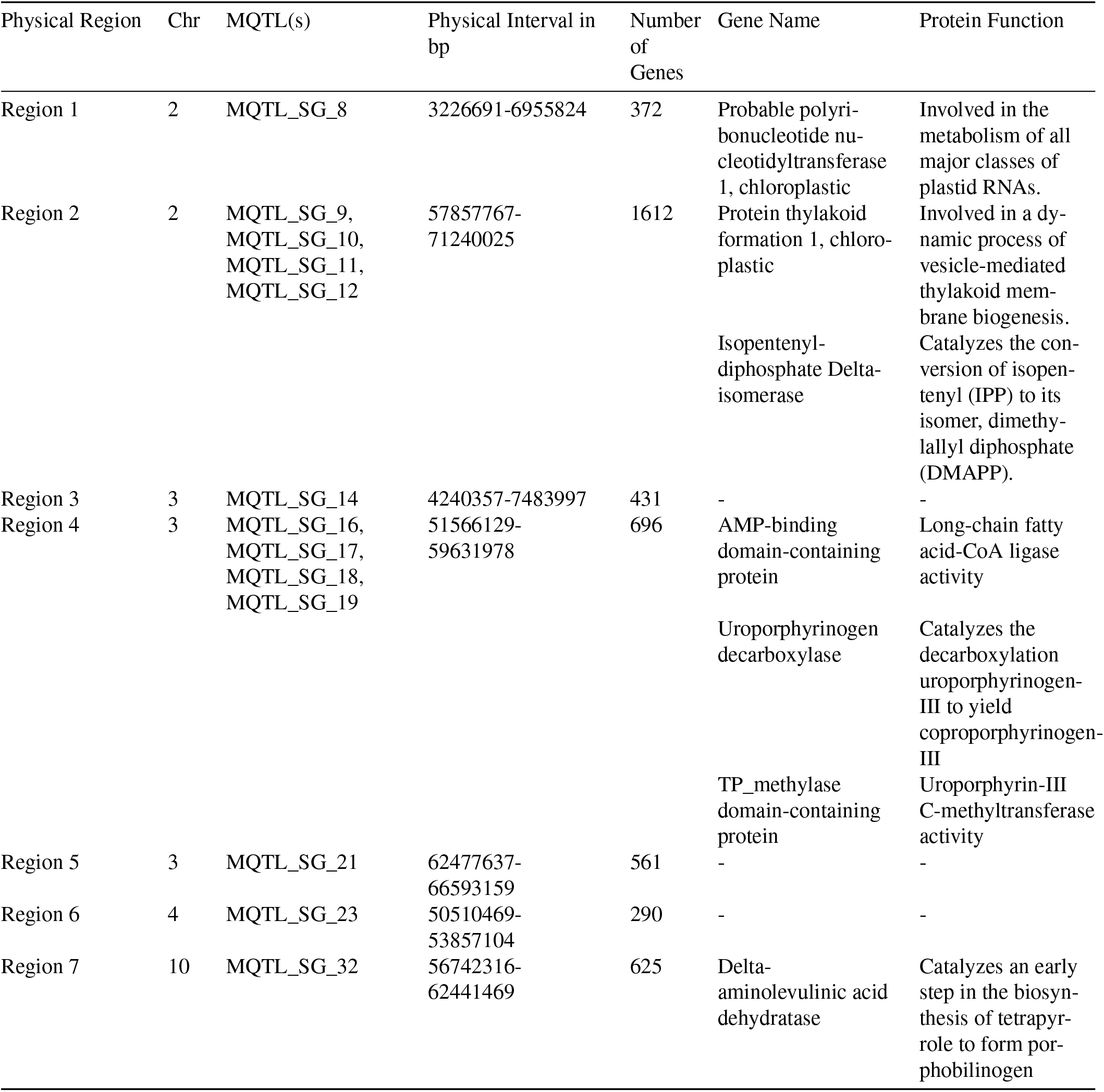
The candidate genes in the MQTL regions and their functions

## Discussion

Though sorghum possesses a tremendous ability to survive droughts, yet it’s yield is sensitive especially at the grain filling stage. One approach for increasing the productivity of crops is to make them resilient to stresses that are detrimental to their productivity. The mechanisms by which crops respond to water deficits are complex, and studies have been able to identify a vast number of traits that govern this response (reviewed in [32]). However, the positive effect of stay-green on drought tolerance in Sorghum has been well established in multiple studies [2], [3], [4]. Stay-green is the ability to resist premature leaf senescence in water limited conditions during the grain filling stage. Plants that stay green during this period tend to maintain their chlorophyll content, thereby alleviating the yield losses during moisture stress [13], [8]. Owing to this, breeders have been breeding for stay-green to indirectly improve drought resilience in Sorghum [35].

Marker-assisted selection or transgenic approaches demand a higher degree of understanding of the genetic control of stay-green for rapid genetic manipulation. Stay-green, like most of the agronomically important traits, is a polygenic trait and therefore has been studied in detail using quantitative genetics and molecular marker technology. Quantitative trait loci (QTLs) for stay-green in sorghum have been located on linkage maps in a variety of genetic backgrounds and environments. The potential of these QTLs has been well substantiated through evaluations using introgression lines. Under terminal drought conditions a positive response of stay-green QTLs on leaf chlorophyll levels and green leaf area has been reported [36]. A positive effect of stay-green QTL has also been observed on plant water use and transpiration efficiency [37]. Grain yield QTL have been reported to be co-localized with stay-green QTL, indicating a strong correlation between the two traits [38].

The present work is motivated by the numerous attempts made to dissect genomic regions for stay-green in sorghum in the last two decades Table 1. 130 QTLs for stay-green in Sorghum were compiled for the present study. And a Meta-QTL analysis was conducted, through which 32 consensus regions where QTLs have been consistently identified were detected. This approach overcomes the constraints of a single QTL association experiment by increasing heterogeneity in populations and environments. It also provides more reliable molecular markers and helps in gene identification. The meta-QTL analysis was done using Biomercator V4.2 which contains the softwares from the MetaQTL package [20], [19]. The analysis included construction of consensus maps, QTL projection and QTL clustering. A summary of the MQTL regions identified in this study is shown in Table 2. The results from the meta analysis also align with the overview index of the QTLs plotted in Fig 2. The MQTLs with higher number of underlying QTLs and also distinctness in experiments reporting the underlying QTLs are thought of as more stable through genetic backgrounds and environments [39]. By this criteria MQTL_SG_3, MQTL_SG_9, MQTL_SG_10, MQTL_SG_18, MQTL_SG_19 can be selected as the most stable MQTLs found in this study. Whereas, MQTL_SG_30 on chr 9 is the most refined MQTL, i.e. it has the smallest confidence interval on the genetic map.

**Fig. 2.**
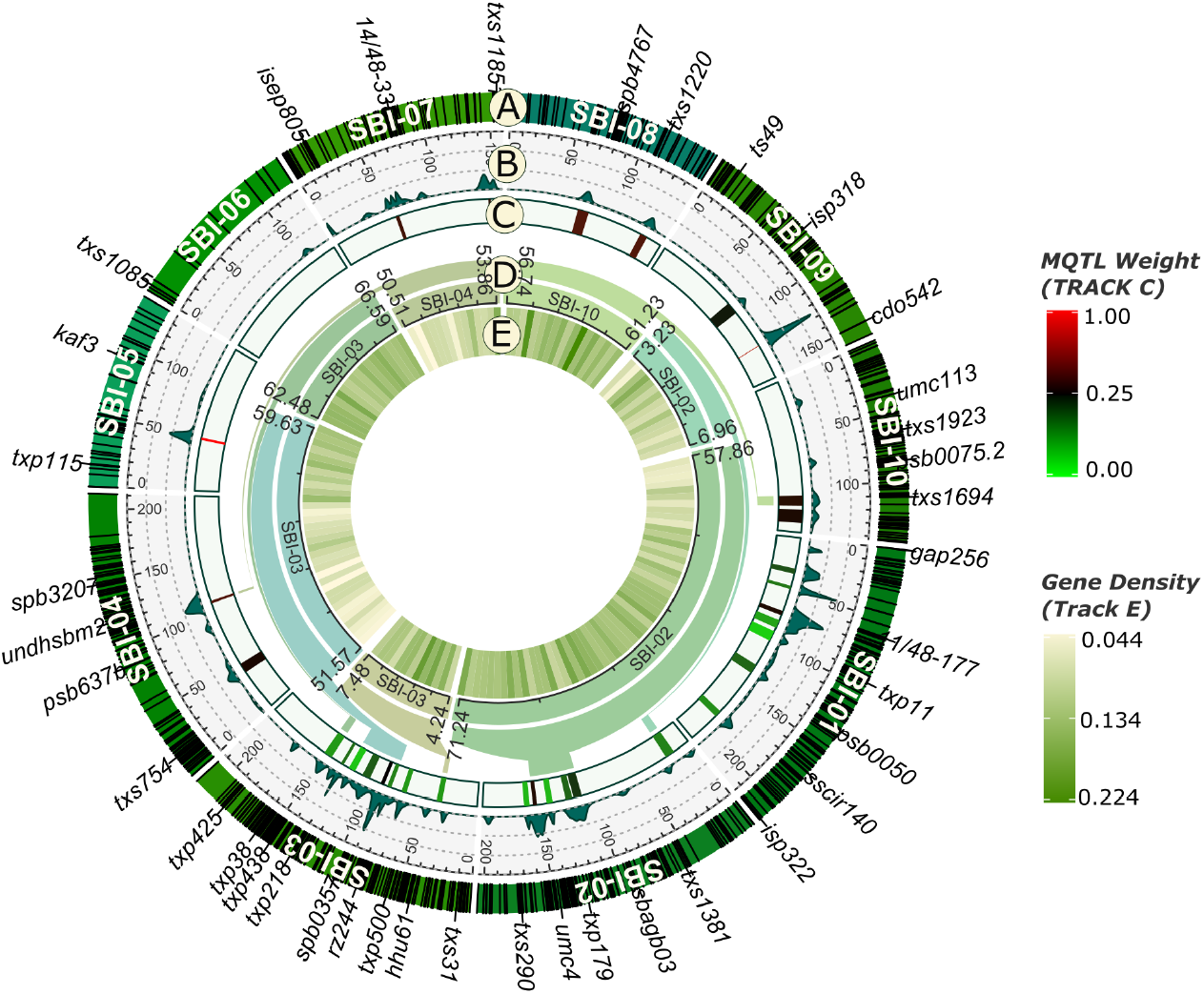
Circular plot showing A) The distribution of genetic markers on the consensus map of Sorghum (In Centimorgan scale); B) Overview index of the original stay-green QTLs; C) Heat map of Meta-QTLs identifies in this study with MQTL weights indicated in the color scale; D) Zooming of MQTLs into the redundant physical interval (in Mega Base-pairs); E) Gene density of the MQTL regions.

The complete genome of sorghum has been sequenced, annotated and published [26]. However, QTLs and MQTLs are genetic intervals, their positions are mapped on linkage maps. And to exactly locate them on the sequenced genome is near to impossible. But an approximation of these intervals is possible through the help of the nearby genetic markers which have been located on the genome. Previous in-silico efforts have been able to map hundreds of genetic markers on the Sorghum genome [21], [22]. But a comprehensive data of genetic markers and their physical location is not available due to unavailability of sequences and limitations of methodologies used to map them. Even with the knowledge of physical location of markers, the exact interval cannot be determined because of the disproportionate relationship between genetic and physical distances. Nevertheless, in the current study we have assumed an approximately linear relationship between distances of genetic markers and physical genomic distances in a local region. And used linear regression to predict the MQTL intervals on the sequenced sorghum genome with 95% confidence level. But the method was successful in predicting a location within the 10 Mb range for only 13 MQTLs. The locations where physical intervals of nearby MQTLs overlapped were reduced to non-redundant regions. Through this 7 regions containing 1-4 MQTLs were obtained Table 3.

Identifying the underlying genes is the ultimate step for any QTL identification study. The CGs in this study were identified in the non-redundant regions on the basis of Gene Ontology terms (GO Biological Process) [40], [31]. All the genes related to either chlorophyll biosynthesis or chlorophyll degradation process within the regions consisting of MQTLs were designated as CGs. The function of orthologous rice genes was acquired from the uniprot database [34]. Two most relevant genes within the MQTL boundaries were Delta-aminolevulinic acid dehydratase (ALA Dehydratase) and Uroporphyrinogen decarboxylase (UPG III Decarboxylase). ALA Dehydratase and UPG III Decarboxylase function closely in the biosynthesis of protoporphyrin IX, a precursor of both hemoglobin and chlorophyll. Another CG, Isopentenyl-diphosphate Delta-isomerase (IPP Isomerase) codes for an enzyme involved in Mevalonate pathway which catalyses the conversion of Isopentyl-diphosphate (IPP) to Dimethyl-allyl diphosphate (DMAPP). During chlorophyll biosynthesis DMAPP is converted to an intermediate geranylgeranyl pyrophosphate, a precursor of chlorophyll. Polynucleotide phosphorylase 1, a gene from MQTL_SG_8, is involved in processing, maturation or degradation of mRNA, tRNA and rRNA of plastids. The probable function for CG coding for TP-methylase domain containing protein is the catalysis of a reaction in which UPG-III is converted to precorrin-2. Two other CGs were functionally involved in thylakoid biogenesis and long chain fatty acid-CoA ligation.

This study was aimed to find relevant genetic intervals that regulate stay-green in sorghum and thereby effect drought resilience. Results from this study could aid in future attempts aiming to improve drought tolerance in Sorghum. Moreover, the information about the genomic regions and the candidate genes will be valuable in understanding the mechanisms of drought response and tolerance. The study was completed using in-silico methods only. Further investigations can focus on fine mapping and validation of the M QTL l oci and markers reported in this study. In the present study CGs were identified using gene ontology annotations, and only genes related to chlorophyll metabolism were selected. The gene sets can be further evaluated through transcriptome analysis to find other genes possibly affecting stay-green.

## Notes

### Competing Interest Statement

The authors have declared no competing interest.

